# Establishing a Canine Urothelial Organoid Repository: A Platform for Comparative and Translational Carcinoma Studies

**DOI:** 10.1101/2025.09.27.678962

**Authors:** Christopher Zdyrski, Aleksandra Pawlak, Hannah F. Nicholson, Megan P. Corbett, Michael Catucci, John Cheville, Jiayi Peng, Corey Saba, Hayden Hamsher, Steven G. Friedenberg, Andrew P. Woodward, Eugene Douglass, Jonathan P. Mochel, Karin Allenspach

## Abstract

Despite the increasing number of treatment options for patients with muscle-invasive bladder Cancer (MIBC), many of these patients ultimately have a poor prognosis. Drug responses vary considerably among patients and in many who respond initially, drug resistance may ultimately develop leading to tumor progression. Scope for improvement has been limited by the phenotypic and molecular diversity of MIBC, which impacts the selection pressures of therapy. To address this translational gap, we describe the largest known canine urothelial carcinoma organoid bioarchive, with in-depth phenotypical and molecular characterization. Immunohistochemistry confirmed the expression of key biomarkers including UPKIII, E-cadherin, Vimentin, and Ki-67 in organoids. Single-nucleus RNA sequencing revealed cellular heterogeneity, while bulk RNA sequencing showed canine patient-to-patient variability in transcriptomic profiling. Bulk RNA sequencing also displayed highly similar expression profiles between tissue- and urine-derived organoids, as well as similarity to previously published human MIBC transcriptome data. Whole genome sequencing in a subset of patients further supported the overall genomic fidelity of the organoids with their tissue of origin. Finally, for proof-of-concept of usefulness for drug testing studies, organoid cytotoxicity to vinblastine was investigated in several organoid lines. These canine bladder cancer organoid lines can be thawed, expanded, and screened for response to potential novel therapeutics to expand personalized medicine approaches. In addition, the organoid lines can serve as a valuable resource for comparative bladder cancer research, and as a pre-clinical screening tool to identify efficacious drugs before taking them into canine clinical trials to support future human clinical trials.

## INTRODUCTION

Bladder cancer (BC) in humans is the most prevalent malignancy of the urinary tract, with approximately 500,000 new cases diagnosed annually worldwide (Ferlay et al., 2021). It is a clinically and biologically diverse disease, broadly divided into non–muscle-invasive bladder cancer (NMIBC) and muscle-invasive bladder cancer (MIBC) (Cheng et al., 2012). About 75% of cases are NMIBC which are those where lesions are confined to the mucosa or submucosa, whereas MIBC invades into or beyond the muscularis propria (Witjes et al., 2021).

For NMIBC, transurethral resection of the bladder tumor (TURBT) can be curative, but MIBC requires multimodal treatment and has a poorer prognosis (Artiles Medina et al., 2025; Saito et al., 2025). Despite neoadjuvant platinum-based chemotherapy followed by radical cystectomy, approximately 50% of patients with MIBC ultimately develop distant metastases, which limit therapeutic options and low survivorship (Krajewski et al., 2021). Newer recommendations include the use of immune checkpoint-inhibitors, with the European Society for Medical Oncology (ESMO) currently recommending a combination of antibody-drug conjugates and immune checkpoint inhibition with or without neoadjuvant chemotherapy for advanced or metastatic disease (Powles et al., 2022). Despite these therapeutic advances, the overall response rate remains low (∼25-40%) (Huang et al., 2025). The use of novel *in vitro* models and the repurposing of already approved oncology drugs could therefore present valuable approaches to accelerate the availability of more personalized treatment options for patients. The most important factor contributing to poor treatment outcomes is believed to be the phenotypic and molecular heterogeneity of MIBC tumors, which limits the translational value of current *in vitro* and *in vivo* pre-clinical models to evaluate therapeutic drug efficacy. Spontaneously occurring invasive urothelial carcinoma (UC) in dogs accounts for approximately 2% of all canine cancers. With an estimated 4–6 million pet dogs diagnosed with cancer annually in the United States, canine UC likely affects over 60,000 animals per year, which is higher than the estimated incidence of MIBC in humans (approximately 20,000 new cases per year in the US) (Gore et al., 2024; Knapp et al., 2014). At the time of diagnosis, most have high-grade histopathologic features and muscle invasion. Approximately 20% of dogs also have distant metastases at diagnosis, which is generally incurable (Knapp et al., 2014, 2020; Mutsaers et al., 2003). Notably, canine UC closely resembles human MIBC in terms of histopathology, molecular features, biological behavior, metastatic patterns, therapeutic response, and clinical outcome (Knapp et al., 2000). These similarities underscore the value of the dog as a comparative model for invasive bladder cancer, including therapeutic evaluation. As companion animals, dogs are increasingly affected by “civilization diseases,” including cancer (Pawlak et al., 2013). Their large body size, shorter lifespan, spontaneous tumor development, and comparable responses to cytotoxic agents make them a valuable translational model, particularly for pre-clinical efficacy assessments (Momozawa, 2019).

Traditional two-dimensional (2D) *in vitro* cancer models have long been used for drug screening, but they fail to replicate the complex architecture, cellular heterogeneity, and tumor– immune interactions characteristic of *in vivo* tumors (Allenspach et al., 2023). Three-dimensional organoid cultures, in contrast, can more accurately mimic cell-to-cell and cell-to-matrix interactions, cell polarity, and the metabolic environment in tumors (Shamir and Ewald, 2014; Whyard et al., 2020; Xia et al., 2019). Such tissue-derived organoids are genetically stable in long-term culture and have been shown to better predict *in vivo* chemosensitivity profiles, particularly in tumors exhibiting vast molecular and phenotypic heterogeneity, such as MIBC (Lee et al., 2018; Vlachogiannis et al., 2018).

Although several groups have reported the establishment of human and mouse bladder cancer organoid bioarchives, these organoid lines are not yet widely available (Burgués et al., 2007; Lee et al., 2018; Mullenders et al., 2019; Santos et al., 2019; Yoshida et al., 2015). Several canine epithelial organoid models have also been established and extensively characterized, supporting their translational value based on conserved molecular and histologic features between the two species (Inglebert et al., 2022; Sato et al., 2023; Scheemaeker et al., 2023; Shiota et al., 2023). These *in vitro* models may provide critical pre-clinical data, which could accelerate clinical trial readiness in humans with MIBC.

Canine UC organoids can be cultured from urine samples or tumor biopsies and exhibit urothelial marker expression and histoarchitecture consistent with invasive UC in dogs and humans (Elbadawy et al., 2019, 2021). Nonetheless, current reports describe only a limited number of canine UC organoid lines, and lack standardization in culture method and molecular characterization, which hampers their utility for broader research applications.

The present study aimed to establish and systematically characterize a larger bioarchive of urine- and tissue-derived canine UC organoids currently catalogued in the Integrated Canine Data Commons of the National Cancer Institute (https://caninecommons.cancer.gov/#/study/ORGANOIDS01). We employed immunohistochemistry, bulk and single-nucleus RNA sequencing, and whole genome sequencing to assess fidelity to the original tumors and to available data on human MIBC, to assess their suitability as a translational preclinical model for therapeutic screening and precision medicine applications in both veterinary and human oncology.

## RESULTS

### Organoid culture and morphology characteristics

A total of 27 organoid lines from 14 dogs were successfully established (patient metadata is presented in **Table 1**), three of these (UC-P1, UC-P2, and UC-P3) were briefly previously described by our group with minimal characterization (Elbadawy et al., 2025). From nine patients, only urine-derived organoids were established. From two patients (P6 and P7), in addition to urine, we were able to establish tissue-derived organoid lines, with P7 being sampled from two sites. From three patients (12, 13, 14), only tissue-derived organoids were created, with P14 having two sample sites. From P6, longitudinal sampling was performed, resulting in the establishment of nine urine-derived organoid lines, in addition to two tissue-derived organoid lines (**Supplemental Figure 1**).

**Table 1.** Patient metadata including patient unique identifier, sampling location, breed, sex, and age. Female spayed = FS, Female intact = FI, Male neutered = MN.

| Unique Patient ID | Organoid Sample Site | Lines Derived | Breed | Age (yr) | Sex |
| --- | --- | --- | --- | --- | --- |
| UC-P1 | Urine | 1 | German shorthaired pointer | 11 | MN |
| UC-P2 | Urine | 1 | blue heeler | 12 | MN |
| UC-P3 | Urine | 1 | mixed breed | 13 | FS |
| UC-P4 | Urine | 1 | mixed breed | 9 | MN |
| UC-P5 | Urine | 1 | beagle | 13 | FS |
| UC-P6 | Urine/Tissue | 9 (urine)<br>2 (tissue) | coonhound mix | 7 | FS |
| UC-P7 | Urine/Mid Bladder/Urethra | 3 | mixed breed | 12 | FS |
| UC-P8 | Urine | 1 | mixed breed | 11 | FS |
| UC-P9 | Urine | 1 | black Labrador retriever | 11 | MN |
| UC-P10 | Urine | 1 | beagle mix | 14 | MN |
| UC-P11 | Urine | 1 | pitbull mix | 10 | FS |
| UC-P12 | Tissue | 1 | Welsh springer spaniel | 12 | MN |
| UC-P13 | Urethra | 1 | German shepherd dog | 8.9 | FS |
| UC-P14 | Apex/Trigone Tissue | 2 | Pomeranian | 13 | FI |

Canine UC organoids typically developed within five days after seeding and were identified as solid or cystic structures. Urine-derived organoids were typically of very high density within the first week, while tissue-derived organoids would grow and bud off the tissue fragments until the first passage and were then released from the anchoring tissue. Canine UC organoid cultures could be established with around 80% efficiency in our laboratory from both urine and tissue and could be propagated for extended periods (over one year), as tested on multiple organoid lines.

Once the cultures were expanded, no notable differences were observed under brightfield microscopic observation regarding their morphology and growth rate between tissue- or urine-derived cell lines. Brightfield images of the formed organoids are shown in **Figure 1A-C** with slight variations between patients and time since passage, with some resembling more cystic structures (i.e. P4 and P8) and others resembling budding structures (i.e. P2 and P3). Formalin-fixed, paraffin-embedded organoids had three main morphologies: solid, porous, or with a larger central lumen, replicating canine UC heterogeneity (**Figure 1A-C**). Predominantly solid organoids were most common, as were organoids that were fenestrated, characterized by variably shaped lumina and microcysts often with basophilic mucoid material present inside the organoids. Some samples had organoids with lumina lined by one or more layers of cuboidal cells, forming tubule-like arrangements. Squamous differentiation, cell swelling, and elongation of the cells were also common findings, and these changes were found in individual organoids from the same sample rather than being a diffuse change in every organoid in any given sample. The cells were polygonal, consistent with epithelial cells, and had varying amounts of eosinophilic cytoplasm. There was cytologic evidence of malignancy in all organoid lines, characterized by moderate to marked variation in cell size (anisocytosis) and nuclear size (anisokaryosis), nuclear crowding, open, clumped chromatin, and prominent, often multiple nucleoli. Additionally, some samples had atypical mitotic figures and signet ring cells, the latter of which has often been associated with carcinomas in previous publications (Meuten and Meuten, 2016). Melamed-Wolinska bodies, considered a cytologic hallmark of urothelial cells (Gordillo and Jiménez-Heffernan, 2022; MELAMED and WOLINSKA, 1961), were rare and only found in four organoid lines. The organization of cells within organoids was not random, and organoids reproduced structures resembling canine invasive UC (**Figure 1C**).

**Figure 1:**
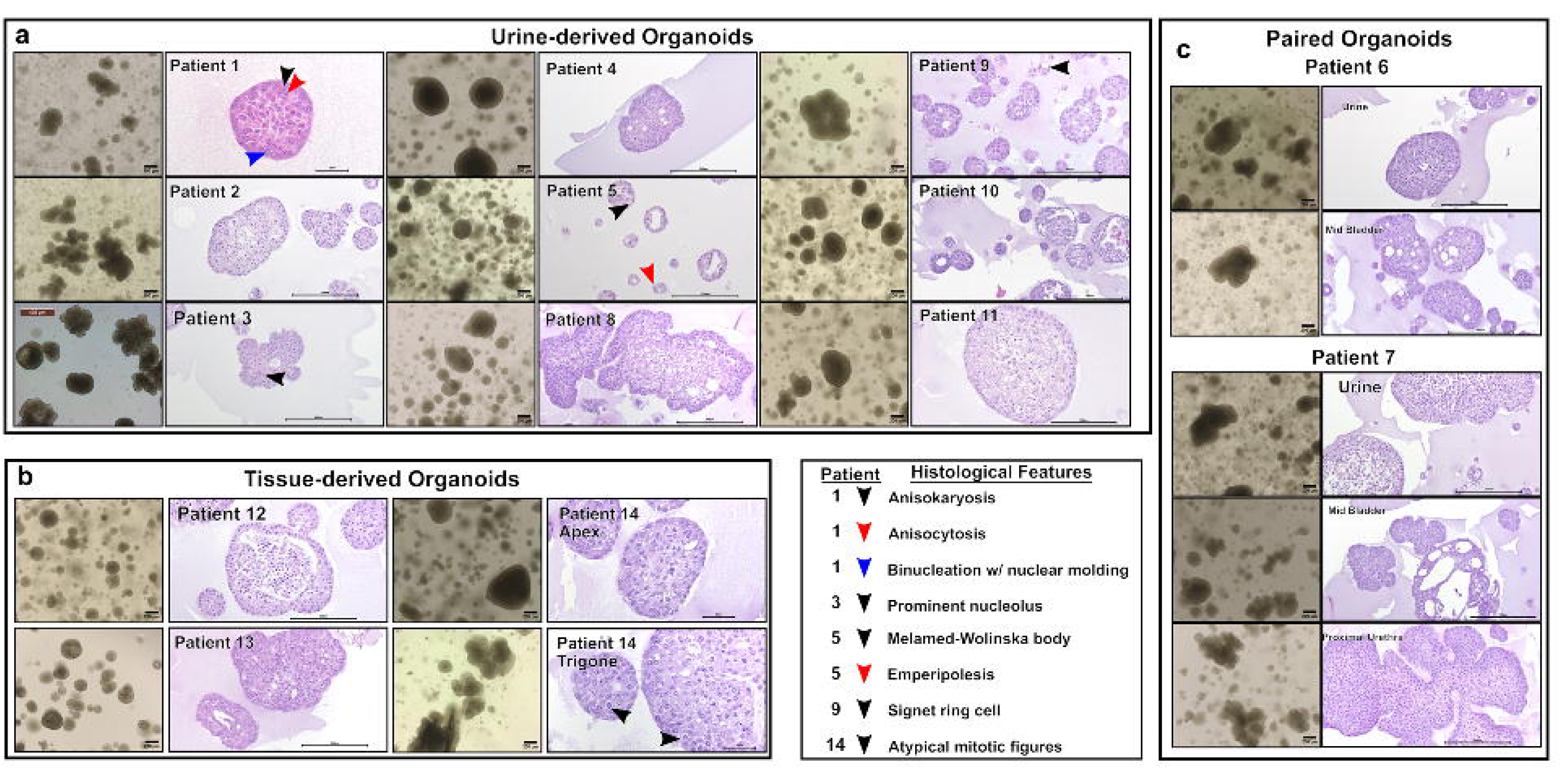
Examples of the morphological and histological phenotypes that can be found in (**a**) urine-derived (**b**) tissue-derived, and (**c**) matched urine-derived and tissue-derived canine UC organoids from the same patients. Notable and representative histological features are denoted with arrows; arrow meanings are denoted in the key. Scale bars are in µm.

Characterization of cell types in the organoid culture was performed with immunohistochemistry (IHC) where paraffin-embedding of the organoid lines was possible. Umbrella cells were identified using uroplakin III (UPKIII) (**Figure 2**), with most organoid lines being immunonegative (11/19 lines) or faintly immunopositive (6/19 lines). Only occasionally was moderate immunopositivity observed (2/19 lines). None of the organoid lines had strong UPKIII expression. The epithelial origin of the organoids was confirmed with E-cadherin, which was, as expected (Jing et al., 2023), mostly strongly positive in the organoid lines (negative in 4/21, faint in 1/21, moderate in 1/21, strong in 14/21, and variable staining in 1/21). Vimentin staining was used to identify mesenchymal-like cells and was variably expressed in the organoid lines (negative in 2/20, faint in 6/20, moderate in 8/20 and strong in 4/20 organoid lines) (**Figure 2**). Ki-67 staining was used to assess proliferation of the organoid lines, with 18/19 organoid lines immunopositive at varying levels (**Figure 2**).

**Figure 2:**
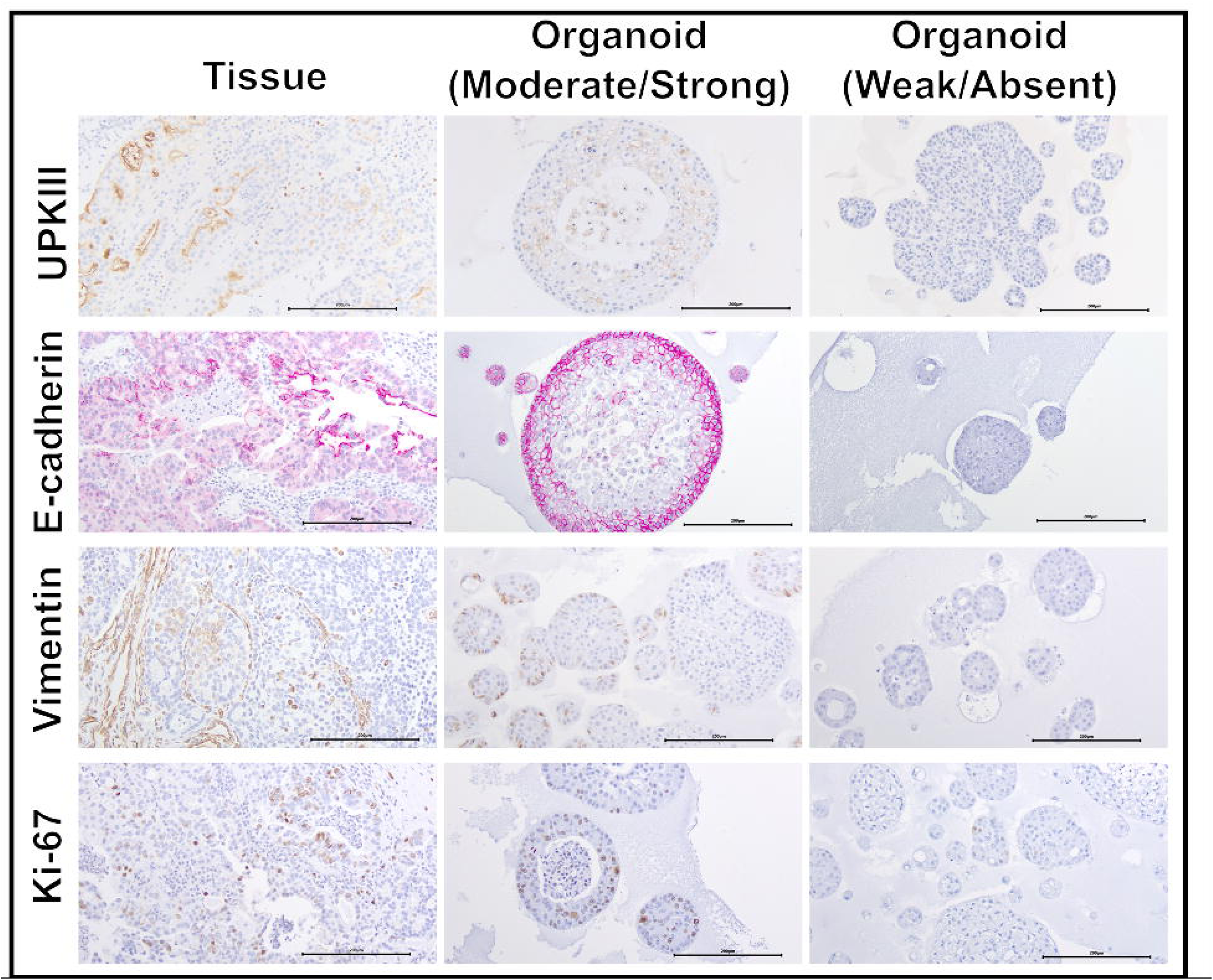
Immunohistochemical appearance of canine UC organoids and corresponding tissues. IHC markers included UPKIII, E-cadherin, Vimentin, and Ki-67. Representative patients were used to demonstrate the variability of Moderate/Strong and Weak/Absent staining across organoids and tissues. Scale bars are in µm.

### Transcriptomic analysis

RNA sequencing analysis was used to define the transcriptomic profile of cells forming organoids in comparison with their corresponding tissue samples, where available, to assess the degree of heterogeneity between individual patients. In addition, we also analyzed selected signaling pathways activated in the organoid lines to identify possible molecular targets that could be therapeutically exploited.

First, principal component analysis (PCA) showed one cluster of all UC organoid lines separate from two distinct clusters from the tissue samples (PC1: 33%), even when comparing paired tissue/organoid samples from the same dog (**Figure 3A**). The two tissue clusters were characterized by enrichment of genes associated with either luminal or basal molecular subtypes, as described in the human MIBC TCGA samples and recent consensus MIBC classifiers across samples (Kamoun et al., 2020; Robertson et al., 2017). In contrast, all organoids exhibited strong enrichment of luminal signatures. These luminal and basal molecular subtypes were confirmed using the consensus MIBC R package (Kamoun et al., 2020) and visualized using cyan and dark red color coding, respectively. When compiling a PCA of organoid lines only, transcriptional differences between organoids were driven by patient-specific differences, highlighting the heterogeneity of canine UC, which is mirrored in this dataset (**Supplemental Figure 2A**). Hierarchical clustering showed that tissues clustered separately from organoids, as expected, due to gene expression associated with cell types present in tissues (fibroblasts, vasculature, immune cells, etc.) that were absent in organoids. The clustering of basal and luminal MIBC was highly correlated between our canine- and human-MIBC samples as illustrated in the correlation matrix in **Figure 3B**, which is annotated with basal and luminal molecular subtypes, based on the human consensus MIBC algorithm (Kamoun et al., 2020). This provides evidence that molecular subtypes of MIBC are conserved across species.

**Figure 3:**
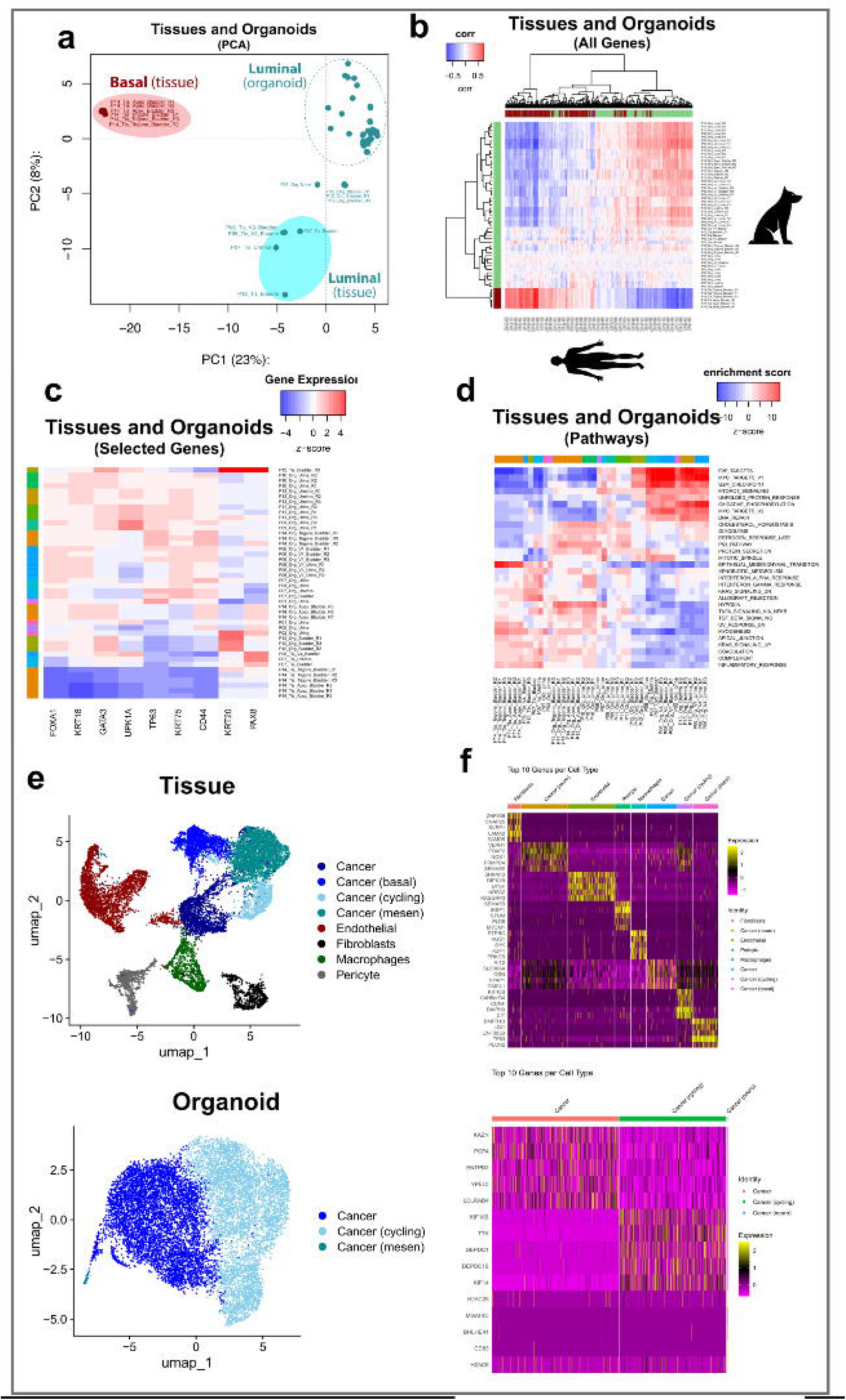
(**a**) PCA showing the distribution of transcriptomes of organoid lines and tissues within the context of luminal (green) or basal (red) phenotypes according to the TCGA classification (Kamoun et al., 2020). (**b**) A heatmap of all gene expression patterns across canine organoids and tissue samples compared to transcriptomic data publicly available from human MIBC tissue samples (Robertson et al., 2017). (**c**) A heatmap displaying the expression profiles of selected genes that are relevant in UC across sample types. (**d**) Pathway analysis for hallmark gene sets across all samples (Liberzon et al., 2015). (**e**) Identification of cell clusters using snRNA-seq for a matched canine UC organoid line and its parent tissue sample. Annotated UMAPs displaying distinct cell clusters (colored) identified in a paired canine tissue and organoid sample. (**f**) Heatmaps identifying the top 10 markers for each cluster (upregulated = yellow and downregulated = purple, with respect to each other), expressed as average log2Fold Change.

Next, to assess the presence of different cell types forming the organoids, we analyzed the transcriptomic profiles for genes that are typically expressed in urothelial cells. This was done to confirm that cellular identity and the ability of the organoid lines to maintain parent tumor heterogeneity was conserved in the *in vitro* culture conditions (**Figure 3C**). The analysis confirmed the presence of different urothelial cell populations and defined their phenotype as: putative urothelial stem cells (CD44+), differentiated umbrella cells (UPK1A+), luminal cells (CK18+, CK20+; FOXA1+, GATA3+), and basal cells (p63+ and CK5+), as shown in **Figure 3C**. The study showed higher expression of the CD44 marker in organoids compared to most tumor tissues, likely due to the increase in stem cell populations in organoid cultures relative to other cell types. High expression of stem cells in bladder cancer organoids has also previously been observed in human organoid studies (Mullenders et al., 2019). The comparison of transcriptomes from tissue- and urine-derived organoids is highlighted in a volcano plot showing both shared and unique upregulated genes across all organoids (**Supplemental Figure 2B**). After FDR correction, 93.4% of genes showed no statistically significant difference, with 6.6% meeting the Q < 0.05 threshold. This modest proportion is close to the rate expected under the null hypothesis, indicating broad transcriptomic similarity between the two organoid sources. Genes upregulated in urine-derived organoids included CXCL8, S100A12, SLPI, and HRH3, while tissue-derived organoids had expression of CDH26, CNTN3, NOS1, and LRP3.

Finally, we analyzed hallmark pathways for our canine UC organoid lines and their corresponding tumor tissues (**Figure 3D**). Hierarchical clustering of the genes showed a similar gene expression pattern among different patients’ organoids and tissues. In the tissues, a clear activation of pathways related to the inflammatory process (specifically Inflammatory response and TNFA Signaling via NFKB pathways), were detected, likely due to inclusion of immune cells in the tissue samples, while in the organoids, activation of pathways related to cell cycle regulation, E2F targets, MYC targets, G2M checkpoint, and MTOR signaling were dominant, likely due to the organoid’s proliferative nature within an *in vitro* environment.

### Single nuclei RNA sequencing

For one patient (P12), we were able to perform snRNASeq for both the tissue and organoid culture. In the tissue, there were 12,895 total cells detected, 52,849 mean reads per cell, and 2,199 median genes per cell. A total of 8 distinct cell clusters were identified by UMAP; among them were basal UC cells (TP63), endothelial cells, fibroblasts, macrophages, and pericytes (**Figure 3E-F**). In the organoid line, there were 11,585 total cells detected, 67,209 mean reads per cell, and 4,428 median genes per cell. A total of three distinct cell clusters were identified by UMAP, including two that were enriched in mesenchymal genes (mesen), or proliferation genes (cycling). In both the tissue and organoid samples, the cycling clusters had the KIF18B gene highly upregulated, which encodes for Kinesin family member 18B and is strongly involved with mitosis, specifically cell division and chromosome segregation, suggesting the active proliferation of the cells (**Supplemental Figure 2C-D**).

### WGS Analyses

To assess the somatic mutational status of the organoids, we performed WGS of the samples, with the aim to evaluate the number of conserved mutations found in the organoid lines compared to their paired tumor tissue samples.

### Whole Genome Sequencing (WGS) of tissues and organoids for variant detection

When analyzing all genes containing high-moderate- or low-impact variants, the majority were shared between the tumor and organoids (**Figure 4A**). When analyzing specific variants, however, most were found to be exclusive to either the tissue or the organoid samples (**Figure 4B**). In the three donors where matched tissue and organoids were available, the top 25 genes by variant count were tracked along with their predicted biological impact (moderate or high). Importantly, a subset of the top 25 genes by variant count from each patient was retained in the organoids. For example, in P12, 9 of 25 genes were the most frequently mutated when comparing between the tissue-derived organoids and primary tumor. Mutations in genes including KRAS and KHK were among two of the top 25 genes mutated by variant count in both tissues and organoids in our cohort (**Figure 4E**).

**Figure 4:**
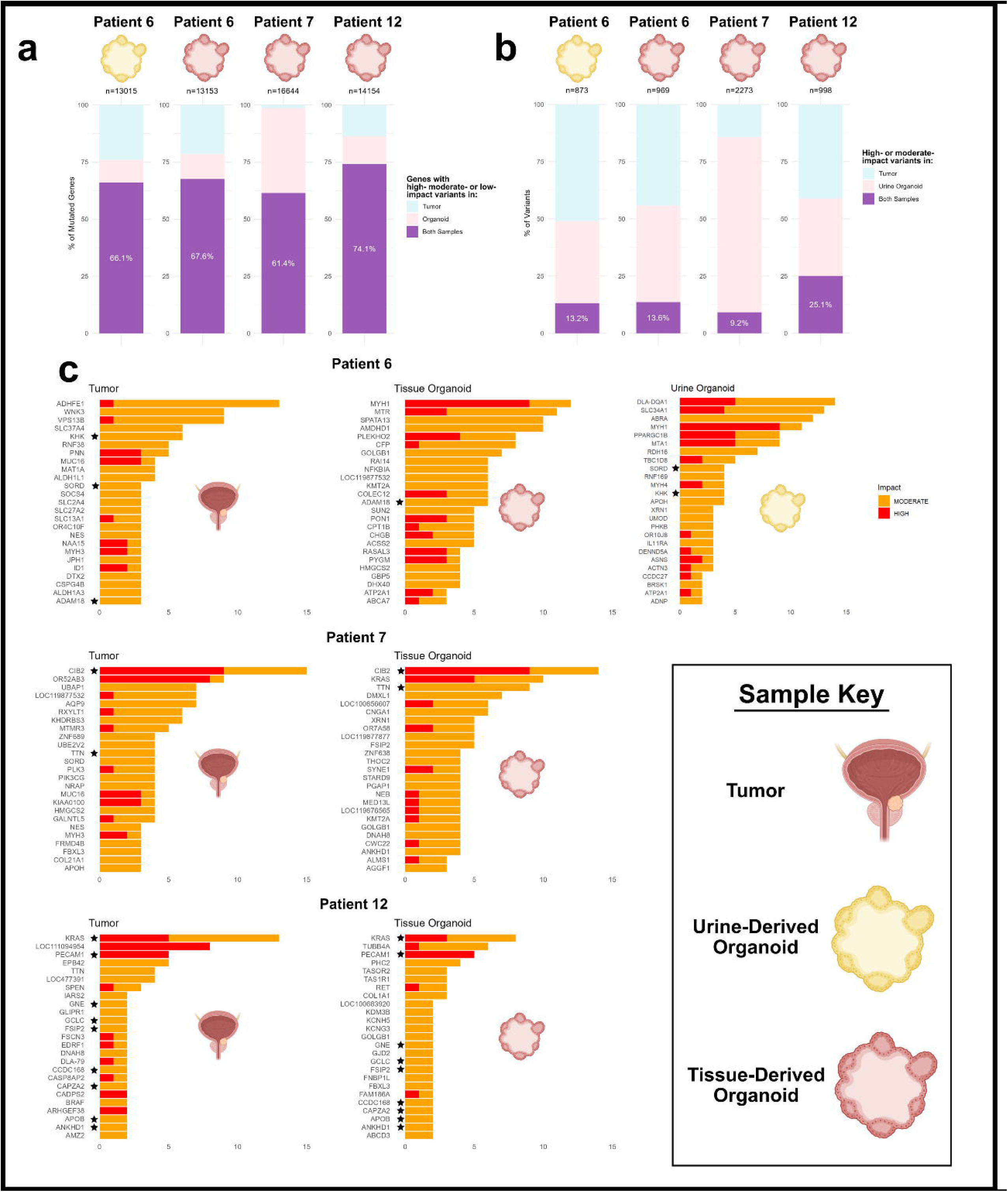
(**a**) Percentage of all genes containing high-moderate- or low-impact variants in both samples (purple), unique to the organoid (pink), or unique to the tumor of origin (blue). (**b**) Percentage of identical high- or moderate-impact variants in both samples (purple), unique to the organoid (pink), or unique to the tumor of origin (blue). (**c**) Top 25 genes by variant count with moderate-(orange) or high-impact (red) variants found in each sample. Stars denote the same gene present in both tumor and organoid. Organoid symbols represent urine-derived organoids (yellow) and tissue-derived organoids (pink). Created using BioRender.com.

BRAF mutations, and particularly, the BRAF deletion mutation V595E (analogous to human BRAF deletion mutation V600E), was evaluated in the dataset as this mutation has previously been identified in >50% of canine bladder cancers (Bartel et al., 2024; Grassinger et al., 2019; Kuo et al., 2024; Thomas et al., 2023). The BRAF deletion mutation was found in the bladder tissues of P6, P7, and P12 in our dataset. However, none of the organoids derived from P6 retained the BRAF mutation. In P7, the urethra-derived organoids retained the BRAF mutation, while the bladder tissue-derived organoids did not. In P12, both tissue and tissue-derived organoids retained the BRAF mutation.

### Cytotoxicity study

To demonstrate the usefulness of UC organoids for drug sensitivity studies, we performed a proof-of-concept cytotoxicity study using vinblastine on two matched urine- and tissue-derived organoids (P6 and P7) (**Figure 5A**). After exposure of the organoid lines to vinblastine, clear morphological changes occurred over the time frame of the 72-hour experiment, with smaller organoids and apoptotic cells appearing over time (**Figure 5B**). The metabolic activity of the cells was measured using an 3-[4,5-dimethylthiazole-2-yl]-2,5-diphenyltetrazolium bromide (MTT) assay, with exposure to vinblastine resulting in a significant decrease in organoid metabolic activity (**Figure 5C**).

**Figure 5:**
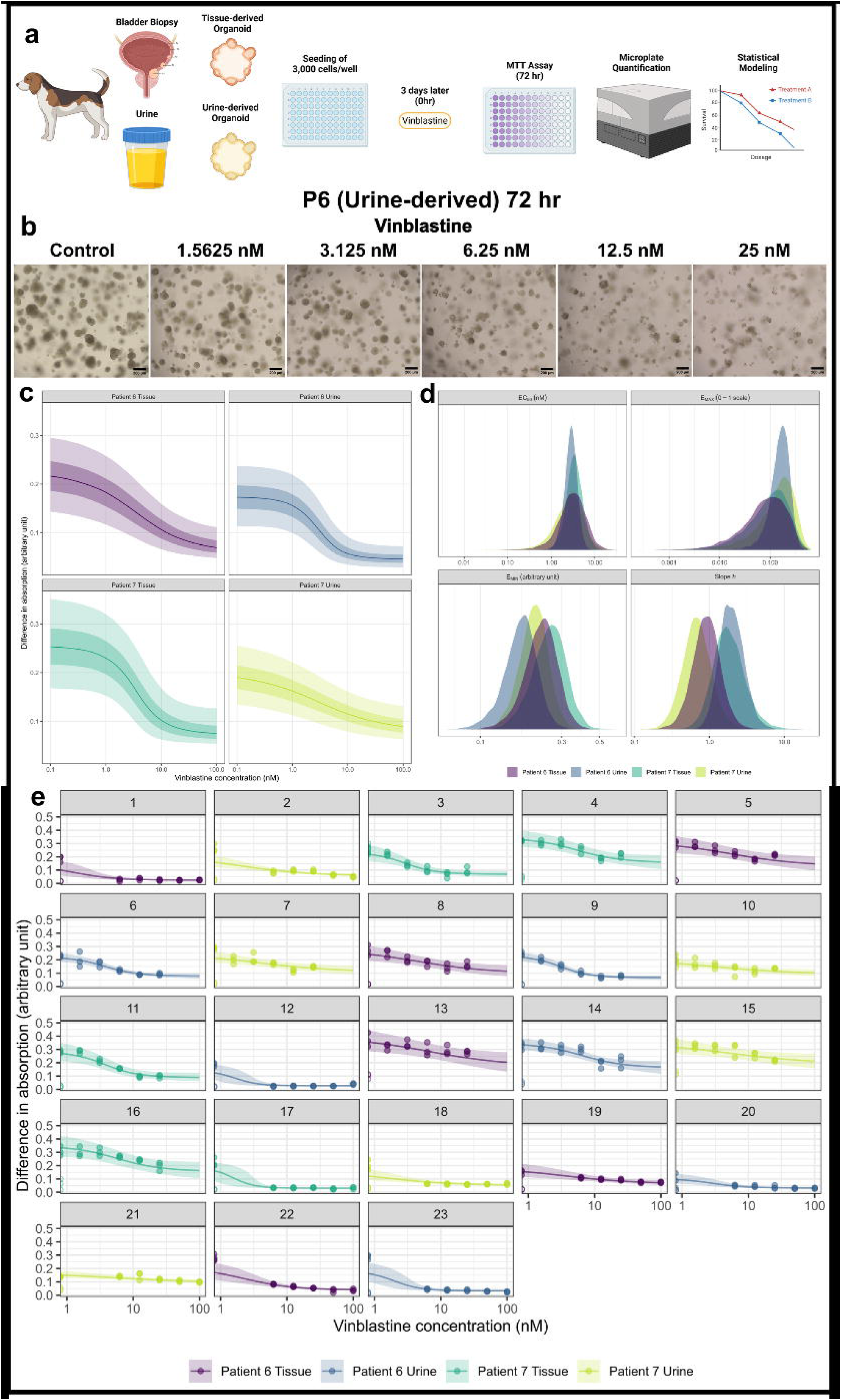
(**a**) Experimental workflow for treatment of urine- and tissue-derived organoids, drug screening assay, and statistical modeling. (**b**) Brightfield images highlighting the morphological changes of controls and vinblastine-treated organoids (controls: no treatment, five concentrations of vinblastine: 1.5625, 3.125, 6.25, 12.5, 25 nM). Scale bars are in µm. (**c**) Posterior-predicted dose-response relationship for organoid viability (MTT assay) by vinblastine concentration (nM), by individual, for a hypothetical average plate (ignoring between-plate variation) The solid line represents the posterior median predicted response, and the fields indicate 50% and 90% credible regions demonstrating uncertainty in the dose-response relationships. (**d**) Posterior probability distributions of the individual pharmacodynamic parameters. (**e**) Posterior-predicted dose-response relationships by plate. Points represent the observed data. Open points are the observations of DMSO-exposed controls, which are taken as the maximum possible inhibitory effect. The solid line represents the posterior median predicted response, and the field the 90% credible region. Panels indicate different plates, and colors indicate different subjects (urine and tissue specimens from two dogs). Created using BioRender.com.

The IC_50_ estimates from the four tested organoid lines are listed in **Table 2**. All patients had low IC_50_ estimates, with vinblastine having a high potency across all patients. Differences between organoids that were either urine or tissue-derived, as well as differences between P6 vs. P7 were minimal across concentrations.

**Table 2.**
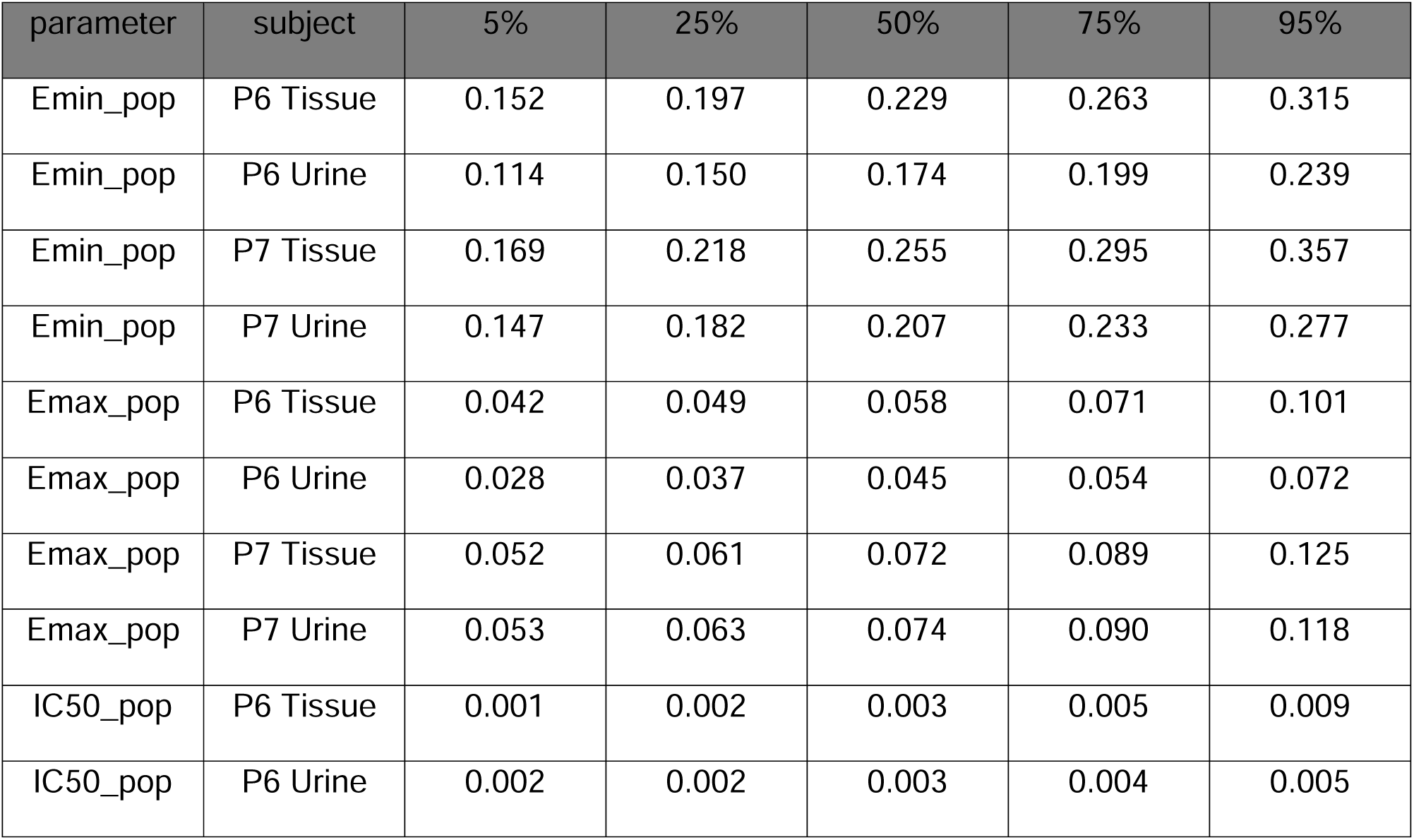
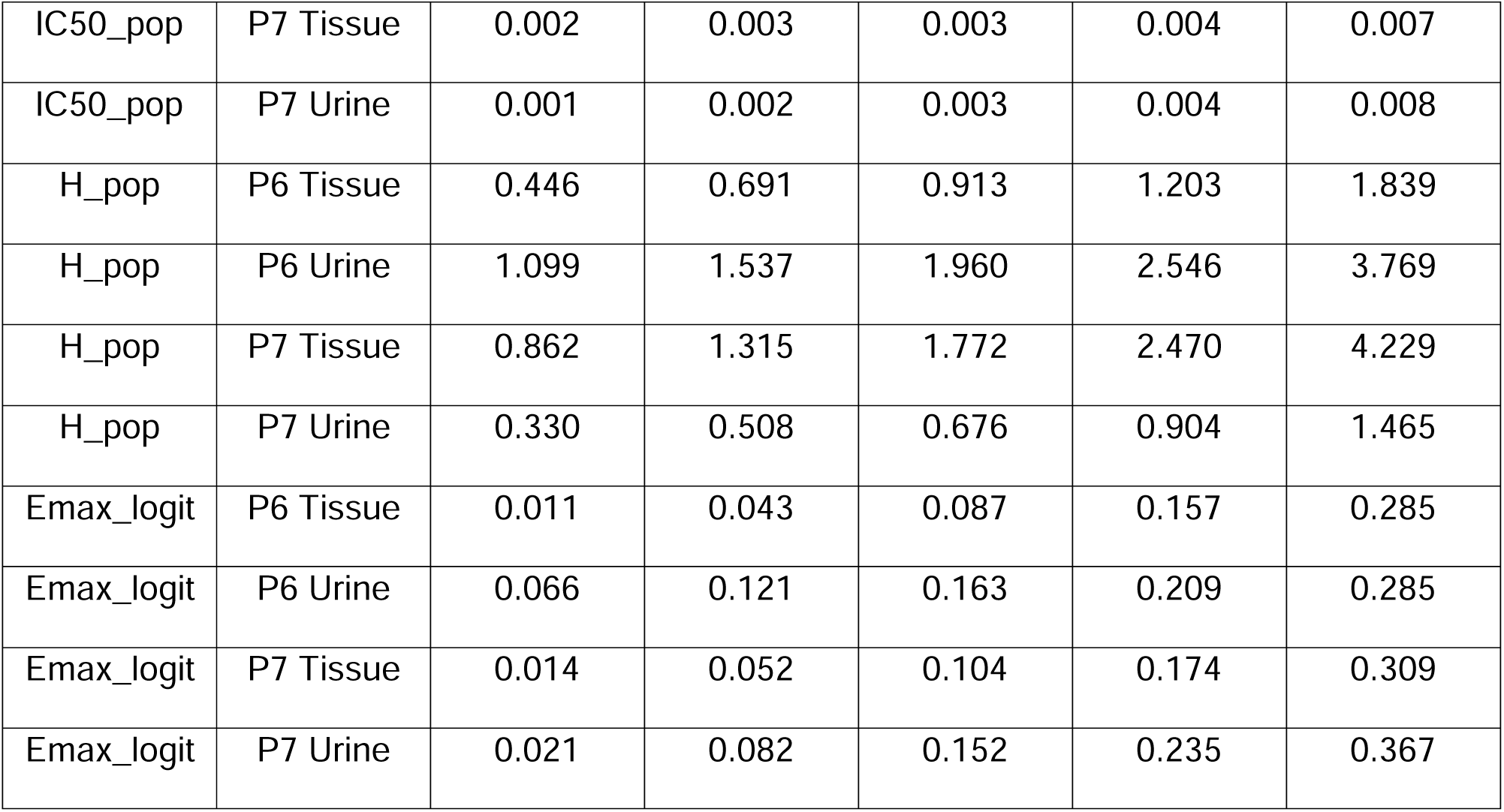
IC_50_ estimates and other parameters given for vinblastine across the two patients for both tissue- and urine-derived organoids.

| parameter | subject | 5% | 25% | 50% | 75% | 95% |
| --- | --- | --- | --- | --- | --- | --- |
| Emin_pop | P6 Tissue | 0.152 | 0.197 | 0.229 | 0.263 | 0.315 |
| Emin_pop | P6 Urine | 0.114 | 0.150 | 0.174 | 0.199 | 0.239 |
| Emin_pop | P7 Tissue | 0.169 | 0.218 | 0.255 | 0.295 | 0.357 |
| Emin_pop | P7 Urine | 0.147 | 0.182 | 0.207 | 0.233 | 0.277 |
| Emax_pop | P6 Tissue | 0.042 | 0.049 | 0.058 | 0.071 | 0.101 |
| Emax_pop | P6 Urine | 0.028 | 0.037 | 0.045 | 0.054 | 0.072 |
| Emax_pop | P7 Tissue | 0.052 | 0.061 | 0.072 | 0.089 | 0.125 |
| Emax_pop | P7 Urine | 0.053 | 0.063 | 0.074 | 0.090 | 0.118 |
| $\text{IC}_{50\_pop}$ | P6 Tissue | 0.001 | 0.002 | 0.003 | 0.005 | 0.009 |
| $\text{IC}_{50\_pop}$ | P6 Urine | 0.002 | 0.002 | 0.003 | 0.004 | 0.005 |
| IC50_pop | P7 Tissue | 0.002 | 0.003 | 0.003 | 0.004 | 0.007 |
| IC50_pop | P7 Urine | 0.001 | 0.002 | 0.003 | 0.004 | 0.008 |
| H_pop | P6 Tissue | 0.446 | 0.691 | 0.913 | 1.203 | 1.839 |
| H_pop | P6 Urine | 1.099 | 1.537 | 1.960 | 2.546 | 3.769 |
| H_pop | P7 Tissue | 0.862 | 1.315 | 1.772 | 2.470 | 4.229 |
| H_pop | P7 Urine | 0.330 | 0.508 | 0.676 | 0.904 | 1.465 |
| E<sub>max</sub>_logit | P6 Tissue | 0.011 | 0.043 | 0.087 | 0.157 | 0.285 |
| E<sub>max</sub>_logit | P6 Urine | 0.066 | 0.121 | 0.163 | 0.209 | 0.285 |
| E<sub>max</sub>_logit | P7 Tissue | 0.014 | 0.052 | 0.104 | 0.174 | 0.309 |
| E<sub>max</sub>_logit | P7 Urine | 0.021 | 0.082 | 0.152 | 0.235 | 0.367 |

## DISCUSSION

The data presented here describes the establishment and in-depth characterization of tissue- and urine-derived urothelial carcinoma (UC) organoids from a large cohort of canine patients, including analyses of their histological features, transcriptomic profiles, mutational landscapes, and responses to cytotoxic agents. This study demonstrates the reproducible generation of canine organoid lines from both tumor tissue and urine samples, supporting their usefulness as an *in vitro* translational model of MIBC.

Initial reports described cancer tissue–originated spheroids (CTOSs) and subsequently patient-derived organoids (PDOs) for human urothelial cancer (Yoshida et al., 2015, 2018), establishing key methods for isolating and culturing bladder cancer organoids. A corresponding study in dogs was published shortly thereafter (Elbadawy et al., 2019), highlighting methodological and phenotypic parallels between species. Since then, several groups have developed protocols for human UC organoid culture (Minoli et al., 2023; Mullenders et al., 2019; Vollmer et al., 2024; Wei et al., 2022), though existing biobanks remain limited and are not widely available (Guo et al., 2024; Radić et al., 2024). Human UC organoid collections typically contain just a few to several dozens of samples, whereas the only previous publication of canine UC organoid cultures remains limited to four lines (Elbadawy et al., 2019), restricting the ability to investigate inter-patient variability in canine bladder cancer. While the first reports of urine-derived UC organoids (urinoids) in dogs focused primarily on establishment of organoid lines (Elbadawy et al., 2019), the analogous method in humans was not described until 2023 (Walz et al., 2023). Although tissue-derived organoids are valuable preclinical models, they are limited in their ability to support longitudinal monitoring or iterative therapeutic evaluation due to the need for repeated invasive sampling. TURBT-derived and cystoscopic biopsy samples are often very small (Abd El-Latif et al., 2013), and are therefore unlikely to represent the actual heterogeneity present within the original tumor. In contrast, urine-derived organoids offer a minimally invasive alternative, though a similar key limitation to tissue biopsies is that only certain cancer stem clones may be included in the organoids, which again limits their ability to reproduce the exact cellular composition of the tumor of origin. Furthermore, the fact that much fewer urinoid lines have so far been published in human medicine may indicate that most human UCs represent NIMBC, which may contain fewer cancer stem cell clones and/or less aggressive cancer stem cells, therefore making it more difficult to culture organoids from urine sediment. Since canine UC represents almost exclusively invasive tumors, data obtained from canine tumor tissue and tumor-derived organoid lines is likely more comparable to human MIBC-derived datasets.

In our study, we had the opportunity to generate both tissue- and urine-derived organoids from a subset of patients and directly compare them with the corresponding tumor tissues. Under similar *in vitro* culture conditions, organoids derived from urine and tissue showed comparable morphological features and growth dynamics. Organoid morphology was largely similar across patients, although certain features, such as Melamed-Wolinska bodies, which are typically observed in UC tissues, were only seen in a subset of cases. Immunohistochemical analysis confirmed UPKIII expression in a small subset of our organoid lines, likely reflecting the loss of differentiated umbrella cells in most bladder cancer tissues, as is typically seen in human UC (Agaimy et al., 2021), or could be due to their stem cell nature and lack of full maturation. Additional markers, including E-cadherin and Vimentin, have also been used to evaluate histological subtypes and possible epithelial–mesenchymal transition of tumor cells in UC, such as sarcomatoid UC, in both humans and dogs (Corbett et al., 2025; Guo et al., 2019). Vimentin expression was variable in our sample set, with most organoid lines staining faintly or moderately. Strong vimentin expression was found in only four samples (P3, P6 from two visits, and P9). Since we did not have paired tissue samples with histological diagnoses in all organoid lines, we are unable to confirm if they were derived from histological subtypes. E-cadherin expression, by contrast, was strongly positive in most cases, consistent with epithelial origin of UC tumors (Singh et al., 2024). Ki-67 staining confirmed proliferative activity in the organoid lines, a feature that has previously been described in human UC-derived organoids (Zhao et al., 2025). In the one organoid line that was Ki-67 immunonegative, this could have been due to that line only having few organoids in the cross section of the paraffin-embedded slide. Marker expression levels varied among patient-derived organoid lines, likely reflecting underlying biological heterogeneity, patient-to-patient variability, time since passage, and/or total passage number. Histopathological evaluation revealed no differences between the two organoid types when derived from the same tumor. Minor differences were observed at the transcriptomic level, however, overall, both urine- and tissue-derived organoids faithfully recapitulated key UC characteristics, with a predominance of luminal transcriptomic characteristics and large populations of proliferative, cycling cancer cells. Similar findings were reported by Viergever et al. (2024), where immunohistochemistry demonstrated highly similar expression patterns among urinoids, tissue-derived organoids, and primary tumors (Viergever et al., 2024).

Bulk and single-nucleus RNA sequencing revealed significant transcriptomic heterogeneity among organoid cultures and across patients. Pathway enrichment analysis underscored the proliferative signature of the organoids relative to primary tumor tissue and confirmed the absence of immune and endothelial cells in our epithelial cultures. Importantly, transcriptomic differences between tissue- and urine-derived organoids were minimal, with the majority of genes shared across both models, potentially indicating that the less-invasive method of sampling (urine) may be sufficient. With regards to specific genes, SLPI, which encodes a protease-inhibitor, was significantly upregulated in our canine urine-derived organoids, consistent with previous findings from Elbadawy et al. (2019), who identified SLPI as the top upregulated gene when comparing urine-derived UC organoids to normal bladder tissue (Elbadawy et al., 2019). Another gene upregulated in our population of urine-derived organoids included CXCL-8 which is also highly upregulated in human UC and is associated with invasiveness and poor prognosis (Bellmunt et al., 2011; Thalita Reis et al., 2012; Wu et al., 2020). Due to the overall luminal organoid signature, further media refinement may enable the transcriptomic signature to be a more basal phenotype if needed (Robertson et al., 2017), and this variability may be due to media, culture conditions, or sample site and patient variability, as the only other canine bladder cancer organoids had higher expression of basal markers than luminal (Elbadawy et al., 2019). Overall, however, the luminal and basal gene signatures found in our dataset were similar to those found in the human TCGA dataset, indicating close resemblance of the canine organoids with molecular subtype identities in human UC (Robertson et al., 2017).

In our study, we used a non-invasive technique of obtaining cancer cells from urine sediment to assess the transcriptome of tumor-derived organoids over the course of the disease and treatments. We obtained urine samples from one patient multiple times over a period of approximately 7 months, including multiple treatments with vinblastine as well as a caninized immune-checkpoint inhibitor treatment. In this longitudinal study, organoid lines were successfully derived from urine samples across all timepoints of sampling, irrespective of treatment modality (e.g., vinblastine-based chemotherapy or immunotherapy) or patient condition. Organoids were established and characterized morphologically and at the transcriptomic level. Although more detailed analyses were limited by sample availability and clinical constraints, these findings demonstrate the feasibility of generating urine-derived organoids (urinoids) longitudinally from the same patient and during therapy. This approach may enable real-time monitoring of disease progression and treatment response, and could serve as a platform for personalized therapeutic decision-making. Longitudinal sampling is a clear advantage of urine-over tissue-derived organoids; however, the tissue-derived organoids may contain additional cancer stem cell clones, possibly explaining the slight variation in transcriptomic and genomic signatures we observed.

Whole genome sequencing (WGS) from three of our canine patients indicated a high concordance between genes mutated in both the organoids and tumor tissue. One human study reported high mutational overlap for UC urinoids and tumor tissues in individual cases (Viergever et al., 2024). Similarly to our study, it was reported that some inter-patient variation existed, with 11 organoid lines showing >80% mutational concordance with their tumor of origin, while 4 lines demonstrated <60% overlap in mutational profiles, highlighting both the potential and the variability inherent to organoid-based modeling (Lee et al., 2018). In general, WGS of our limited dataset demonstrated that the genes being mutated were similar between organoids and the corresponding tumors. However, when comparing the exact variants of high- and moderate-impact in the tumor of origin and organoid lines, discordances were observed, likely due to differences in cell content and hence, cancer stem cell clones, between tissues and organoids. In addition, clonal selection imposed by *in vitro* culture conditions are also possible explanations. While there are scarce numbers of bladder cancer organoid studies with corresponding mutational analysis to the parent tumor, summaries have generally shown high— but not complete—concordance between organoids and their parent tumors (Medle et al., 2022). In our sample population, KRAS mutations were found to be the most mutated gene in the P12 tumor and organoids. KRAS mutations are frequently found in human MIBC and are associated with higher tumor grade/stage as well as worse prognosis (Almassi et al., 2022; Li et al., 2023; Ouerhani and Elgaaied, 2011; Smal et al., 2016). Interestingly, the BRAF deletion mutation V595E (analogous to human BRAF deletion mutation V600E) (Mochizuki et al., 2015) was not found in all of our organoid lines, even among those derived from tissues that did have the BRAF mutation. This could be due to the possibility of clonal selection or the media composition driving certain cell populations within the organoids in *in vitro* settings.

In addition to precision-medicine applications, an equally important use of organoid cultures lies in drug development and pre-clinical screening. Human UC organoids have been increasingly used in moderate-to-high-throughput drug screening platforms to assess their potential to predict clinical drug efficacy (Garioni et al., 2023; Minoli et al., 2023; Seiler et al., 2023). For instance, Minoli et al. demonstrated the clinical relevance of organoids by comparing responses to various chemotherapeutic agents (Minoli et al., 2023). Their results showed broad efficacy of gemcitabine and doxorubicin in NMIBC organoids, consistent with prior studies (Han et al., 2021; Li et al., 2020), while cisplatin showed potent cytotoxicity in MIBC organoid models, reflecting its use in clinical treatment protocols (Minoli et al., 2023). In our study, we evaluated the sensitivity of both tissue- and urine-derived organoids to vinblastine—a chemotherapeutic agent commonly used in canine UC. We also assessed potential differences in chemosensitivity between organoids derived from tissue versus urine. Vinblastine elicited a consistent, strong cytotoxic response across all organoid lines tested, in line with previous clinical reports (Arnold et al., 2011; Kaye et al., 2015) and earlier organoid studies (Elbadawy et al., 2019). Unfortunately, due to various treatments and limited sample numbers, comparisons of *in vivo* patient response and *in vitro* organoid response was not possible in this study, but is planned for future investigations.

Regarding sample origin, urine-derived organoids exhibited slightly greater sensitivity to vinblastine than tissue-derived organoids. Although only two patient-derived lines were tested in this study, the apparent pattern aligns with the inter-patient variability reported in other studies using organoids for drug screening (Zhao et al., 2025). While drug sensitivity testing is a common application for organoids, additional uses such as prediction of immune checkpoint inhibitor treatment response, which requires co-culture with autologous immune cells (Feng et al., 2025; Li et al., 2025; Wang et al., 2025), and mechanistic studies focusing on therapeutic target identification, are gaining traction (Guo et al., 2024; Radić et al., 2024). It is important to note that methodological differences across studies using tissue-derived organoids—such as the exact sample source, media composition, plate coating, extracellular matrix type, culture duration, and expansion protocols—can impact cross-study comparisons of transcriptomic and phenotypic data (Guo et al., 2024; Medle et al., 2022; Neuhaus et al., 2023; Radić et al., 2024). We would therefore like to emphasize the importance of clearly reporting these variables to enhance reproducibility and comparability of such data.

This study is subject to several limitations. The organoids tumorgenicity should be checked in the future using methods such as xenografts to confirm their tumor origin, as the presence of healthy cells is possible in these adult stem cell-derived models. For urine-derived samples, the noninvasive nature of collection precluded access to matched tumor tissue and detailed histopathologic characterization in a majority of cases. For WGS analysis, paired germline DNA samples (e.g., blood or adjacent non-tumorous bladder) were not available, limiting our ability to distinguish somatic from germline variants. Additionally, bulk RNA sequencing, while informative, masks cellular heterogeneity and may obscure the presence of minor cellular populations such as fibroblasts or immune cells, which are critical for modeling tumor microenvironment interactions. While snRNASeq overcomes some of the challenges associated with bulk RNASeq, it was limited to one patient in this study and did not include cytoplasmic RNA. These transcripts can, however, be analyzed using scRNASeq, allowing for a more granular analysis, but requires dissociation of the tissue.

## CONCLUSION

In this study, we further expanded our existing canine organoid biobank catalogued on the ICDC portal, including a total of 14 canine cases and 27 organoid lines. We report a comprehensive characterization of the organoid lines using histological, molecular, and genomic techniques to investigate both the potential and the limitations of this model system. With limited human bladder cancer organoid biobanks available to date, and even fewer from canines, this study serves as a proof-of-concept for personalized medicine applications in veterinary medicine with potential translational capabilities. Organoids can play a role in pre-clinical screening prior to animal and human testing to reduce costs and accelerate clinical trials. Overall, this work provides the groundwork for more comprehensive datasets and standardization in veterinary research, which forms the foundation for additional translational research and applicability in human medicine.

## METHODS

### Sample processing

Samples for organoid culture were obtained from either urine or tissue biopsies of canine patients diagnosed with bladder cancer and, if collected at a different site, were placed in shipping media previously described (Sato et al., 2023), with the current composition being reported more recently (Nicholson et al., 2025), immediately after sample collection. All sample collections derived from canine patients at Iowa State University (IACUC-21-250) and the University of Georgia (A2023 10-002-A1) were used under approved IACUC protocols. All biospecimen collections obtained from Purdue University were conducted under an approved IACUC protocol (#1111000124), and tissue or urine samples were shipped to ISU or UGA. Tissue samples were sectioned, with a subset immediately preserved in RNAlater (Invitrogen; AM7021), while the remaining pieces were submerged in shipping media for transport. Urine samples were collected by either catheter or free catch and spun at 600 x g for 3 minutes at 4°C. The supernatant was discarded and ∼10 mL of shipping media was added to the pellet and shipped to the lab on ice packs.

Upon arrival, urine samples were washed in PBS (Corning; 21-040-CM) and if red blood cells (RBC) were present, RBC lysis (Roche; 11814389001) was performed. After a final washing step, Matrigel (originally Corning; 356231 and later Corning; 356255) was added to the pellet, resuspended, and plated (30 µL per well) in a 24-well culture plate (Corning; 3524). The sample was incubated at 37°C for 20 min before the addition of culture media, which was slightly modified from the first patients to the final composition listed in Nicholson et al. (Nicholson et al., 2025). Tissue samples were minced with a scalpel, washed in Advanced DMEM/F12 (Gibco; 12634010), and then processed similarly as urine samples.

### Organoid culture

Formed organoids were maintained as described elsewhere in detail (Nicholson et al., 2025). Organoids were cleaned or passed until a culture of the desired purity and cell number was obtained. Briefly, organoids were resuspended in Cell Recovery Solution (Corning; 354270), incubated at 4°C for 10 minutes, and spun (100 x g for 5 minutes at 4°C). If passing, organoids were then incubated with TryPLE Express (Gibco; 12604-021) and washed in DMEM/F12 (100 x g for 5 minutes at 4°C) prior to resuspension in fresh Matrigel, plating, and addition of fresh culture media. The expanded organoids were then frozen using CryoStor CS10 (Biolife Solutions; 210102) and stored in liquid nitrogen.

### Fixation and H&E staining

For hematoxylin and eosin (H&E) staining, organoid media was removed, and 500 µL of Formalin-acetic acid-alcohol, (FAA, composition in Gabriel et al. 2022) made from ethanol (VWR: 71002-512), acetic acid (Fisher; A38500), formaldehyde (Fisher; F79P4), and MilliQ water, was added to each well (Gabriel et al., 2022). After 24 hours, FAA was replaced with 70% ethanol, which was made from diluting 100% ethanol, and samples were paraffin-embedded, mounted on slides, and stained at the Iowa State University or the University of Georgia Histopathology laboratory. Tissues were fixed in paraformaldehyde and paraffin-embedded according to standard histology procedures.

### Immunohistochemistry

Slides were cut and stained at the University of Georgia Histopathology core for Vimentin V9 (BioGenex; MU074-UC), E-cadherin (BD Biosciences; 610181), and Ki-67 (Cell Marque; 275R-16). Slides were additionally sent to Cornell University for UPKIII staining (Fitzgerald Industries; 10R-U103ax). Slides were then imaged using an Olympus BX41 microscope with a BioVid 4k camera (LW Scientific). Images were color-corrected, and scale bars were added with ToupView, version x64 131 4.11.23945.20231121 (LW Scientific).

### RNA Extractions

Organoids were preserved for bulk RNA-sequencing by removing them from Matrigel using Cell Recovery Solution, as previously described (Nicholson et al., 2025). The cell pellet was resuspended in 100 µL PBS, moved to a cryovial containing 900 µL RNAlater, and stored at - 80°C. For RNA extractions, organoid samples were thawed and transferred to 15 mL tubes, washed with 2 mL of PBS, and centrifuged at 1,200 x g (4°C) for 5 minutes. Then the supernatant was removed and the cell pellet was resuspended in 1 mL of Trizol (Invitrogen; 15596026). Next, each sample was briefly vortexed. Tissues that were either flash frozen or stored in RNALater (if stored in RNALater a wash with PBS was done) were transferred from cryovials to microcentrifuge tubes, 800 µL of Trizol were added to each tube, and tissues were homogenized using a pestle. Both samples (tissue and organoids) were left at room temperature for five minutes prior to centrifuging at 12,000 x g (4°C) for 10 minutes. Supernatant was then transferred to a microcentrifuge tube, and 160 µL or 200 µL of chloroform (Alfa Aesar; J67241) were added for tissues and organoids, respectively. The samples were then vigorously mixed by shaking for 20 seconds and after sitting at room temperature for 2-3 minutes, the samples were spun at 10,000 x g for 18 minutes (4°C). The aqueous top layer was collected and moved to a sterile RNase-free tube before adding an equal amount of 100% RNA-free ethanol (Fisher; BP2818-500). Up to 700 µL were loaded in a Qiagen RNeasy column and collection tube (RNeasy Mini kit; Qiagen; 74104). Samples were then centrifuged at 8,000 x g for 30 seconds, the collection tube waste was discarded, and DNase treatment was performed according to Qiagen’s protocol (Qiagen; 79254). In a new collection tube, 500 µL buffer RPE was added to the column, and the samples were again spun. After discarding the flow-through, this step was repeated, with samples now spun for 2 minutes at 8,000 x g. The flow-through was discarded again, and the samples were centrifuged at the same settings for 1 minute. Then, 50 µL of RNase-free water (Sigma; W4502-50ML) was added to the column and incubated for 2 minutes. Samples were centrifuged twice at 8,000 x g for 1 minute. RNA concentration was determined with a Nanodrop ND-1000 Spectrophotometer (Thermo Fisher Scientific), and samples were kept at −80°C until shipped.

### Bulk RNA sequencing

Three of the samples, UC-P1, UC-P2, and UC-P3, were prepped and sequenced at the Mayo clinic using the names OR-B, OR-A, and OR-E respectively. All other samples had library preparation and sequencing conducted at Azenta Life Sciences (South Plainfield, NJ, USA) as follows: RNA samples were quantified using Qubit 2.0 Fluorometer (ThermoFisher Scientific, Waltham, MA, USA) and RNA integrity was checked with 4200 TapeStation (Agilent Technologies, Palo Alto, CA, USA). Strand-specific RNA sequencing library was prepared by using NEBNext Ultra II Directional RNA Library Prep Kit for Illumina following manufacturer’s instructions (NEB, Ipswich, MA, USA). Briefly, the enriched RNAs were fragmented for 8 minutes at 94 °C. First strand and second strand cDNA were subsequently synthesized. The second strand of cDNA was marked by incorporating dUTP during the synthesis. cDNA fragments were adenylated at 3’ends, and indexed adapter was ligated to cDNA fragments. Limited cycle PCR was used for library enrichment. The incorporated dUTP in second strand cDNA quenched the amplification of second strand, which helped to preserve the strand specificity. The sequencing library was validated on the Agilent TapeStation (Agilent Technologies, Palo Alto, CA, USA) and quantified by using Qubit 2.0 Fluorometer (ThermoFisher Scientific, Waltham, MA, USA) as well as by quantitative PCR (KAPA Biosystems, Wilmington, MA, USA). The sequencing libraries were multiplexed and clustered onto a flowcell on the Illumina NovaSeq instrument according to manufacturer’s instructions. The samples were sequenced using a 2×150bp Paired End (PE) configuration. Image analysis and base calling were conducted by the NovaSeq Control Software (NCS). Raw sequence data (.bcl files) generated from Illumina NovaSeq was converted into fastq files and de-multiplexed using Illumina bcl2fastq 2.20 software. One mis-match was allowed for index sequence identification.

### Nuclei isolation and cell counting

For single-nuclei samples, frozen tissues were received at Azenta, South Plainfield, NJ, USA in dry-ice and stored in liquid nitrogen until further processing. Nuclei extraction was performed using the Miltenyi Nuclei Extraction Buffer (Miltenyi Biotec, Auburn, CA, USA) following manufacturer’s guidelines with gentle MACS Dissociation and C tubes. Upon isolation, the nuclei were counted using AO/PI dye on the Nexcelom Cellaca MX before loading onto the Chromium Controller.

### 3’ RNA library preparation and sequencing

Single-nuclei RNA libraries were generated using the Chromium Single Cell 3’ kit (10X Genomics, CA, USA). Loading onto the Chromium Controller was performed to target capture of ∼6,000 GEMs per sample for downstream analysis and processed through the Chromium Controller following the standard manufacturer’s specifications. The sequencing libraries were evaluated for quality on the Agilent TapeStation (Agilent Technologies, Palo Alto, CA, USA), and quantified using Qubit 2.0 Fluorometer (Invitrogen, Carlsbad, CA). Libraries were quantified using qPCR (Applied Biosystems, Carlsbad, CA, USA) prior to loading onto an Illumina NovaSeq XPlus instrument. The samples were sequenced at a configuration compatible with the recommended guidelines as outlined by 10X Genomics. Raw sequence data (.bcl files) were converted into fastq files and de-multiplexed using the 10X Genomics’ cellranger mkfastq command.

### Bulk and single-nuclei RNA sequencing analysis

After sequencing, read quality was assessed using FastQC, and high-quality reads were mapped to the canine reference genome CanFam3.1. Kallisto or CellRanger was used to conduct read alignment and gene expression counts for bulk and single-nuclei RNA sequencing, respectively (Bray et al., 2016; Zheng et al., 2017). Next, single-nuclei RNA sequencing data was normalized and log transformed using R package Seurat (Butler et al., 2018; Hao et al., 2021, 2024; Satija et al., 2015; Stuart et al., 2019) R and corresponding R scripts were then used for data preparation procedures, analysis procedures, and visualization (R Core Team, 2018). The data was stored on GitHub and figshare repositories, and Zenodo was used to create Digital Object Identifiers (DOIs) for publication citations. Gene Set Enrichment Analysis with the fgsea package was used for pathway analysis, and gene signatures were acquired from the Molecular Signatures Database (MSigDB) (Korotkevich et al.; Liberzon et al., 2011).

### DNA extractions

Organoids were preserved for whole genome sequencing (WGS) by cleaning as before, washing in PBS, and transferring to a cryovial for a final spin. Dry cell pellet or tissue samples were stored at −80°C. A Qiagen Blood and Tissue Kit was used on both tissues and organoid cultures to extract DNA. The manufacturer’s protocol was followed with the addition of RNase A (Thermo Fisher Ref: EN0531) to degrade excess RNA. A Nanodrop was used to quantify the DNA and samples were immediately stored at −80°C. Samples were then shipped on dry ice to Genewiz for QC, preparation, and sequencing.

### Whole genome sequencing

Genomic DNA was quantified using the Qubit 2.0 Fluorometer (ThermoFisher Scientific, Waltham, MA, USA). NEBNext® Ultra™ DNA Library Prep Kit for Illumina, clustering, and sequencing reagents were used throughout the process following the manufacturer’s recommendations. Briefly, the genomic DNA was fragmented by acoustic shearing with a Covaris S220 instrument. Fragmented DNA was cleaned up and end-repaired. Adapters were ligated after adenylation of the 3’ends followed by enrichment by limited cycle PCR. DNA libraries were validated using a High Sensitivity D1000 ScreenTape on the Agilent TapeStation (Agilent Technologies, Palo Alto, CA, USA) and were quantified using Qubit 2.0 Fluorometer. The DNA libraries were also quantified by real time PCR (Applied Biosystems, Carlsbad, CA, USA). The sequencing libraries were multiplexed and clustered onto a flowcell on the Illumina NovaSeq instrument according to manufacturer’s instructions. The samples were sequenced using a 2×150bp Paired End (PE) configuration. Image analysis and base calling were conducted by the NovaSeq Control Software (NCS). Raw sequence data (.bcl files) generated from Illumina NovaSeq was converted into fastq files and de-multiplexed using Illumina bcl2fastq 2.20 software. One mismatch was allowed for index sequence identification.

### Whole genome analysis

FASTQ files obtained from sequencing were aligned to the UU_Cfam_GSD_1.0 canine reference genome (Wang et al., 2021) using the Whole Animal Genome Sequencing (WAGS) pipeline (Cullen and Friedenberg, 2023) to generate binary alignment maps (BAM files). Each sample’s BAM file was individually genotyped using Mutect2 (Benjamin et al., 2019) in tumor-only mode with a --max-mnp-distance parameter set to zero. The resulting genotype records were merged into a single multi-sample file and normalized using bcftools (Danecek et al., 2021). To remove suspected germline variants, the combined file was filtered with GATK SelectVariants --discordance (Auwera and O’Connor, 2020), excluding all variants present in an internal database of whole genome sequences derived from 3,023 dogs, wolves, and coyotes of 402 diverse breeds. This database includes 1,971 dogs, wolves, and coyotes released by the Dog10K consortium (Meadows et al., 2023). High-confidence variant sites were retained by selecting variants where at least one sample had sequencing depth (DP) > 5, variant allele frequency (AF) > 0.01, and tumor log-odds score (TLOD) > 6.

### Cytotoxicity study

Vinblastine (sulfate) (Cayman Chemical; 11762) was purchased as powder, dissolved in DMSO (Millipore Sigma; D2650-100ML) to prepare a stock solution, and stored at −20°C until use. Organoids derived from two patients (P6 and P7) from both urine and tissue samples were used in the study. The cells were plated at a density of 3,000 cells per well in 10 µl of Matrigel in 96-well plates (Corning; 3596) 3 days before the experiment and then were exposed to increasing concentrations of drug for 72 hours (DMSO concentration never exceeded 0.1%, which is considered harmless to the cells). Cells were plated in 3 replicates (3 wells), and the entire experiment was repeated 3 times. Vinblastine was tested in a concentration range of 1.5 – 100 nM. Cells incubated in the medium alone served as a control, 4% DMSO was used as a death control. After 72 hours, 10 μL of MTT solution (5 mg/mL) (Sigma Aldrich; M2003) was added to each well. After 2 hours, medium with MTT was removed and 50 µl of 2% SDS was added to dissolve Matrigel, for another 1 hour. Then, 100 μL of DMSO was added, and after dissolution of the content, absorbance was measured at 570 nm with a spectrophotometric microplate reader (Spark, Tecan, Maennedorf, Switzerland).

### Pharmacodynamic data analysis

Analysis of the dose-response data was conducted in Stan (Carpenter et al., 2017) via R (R Core Team, 2024) with visualization from packages ‘ggplot2’ (Wickham, 2016) and ‘ggdist’ (Kay, 2024). The structural model was four-parameter log-logistic following (Ritz et al., 2015), allowing for between-subject and between-plate variation in each of the pharmacodynamic parameters E_MIN_, E_MAX_, EC_SO_, and H (slope). Subjects were allowed to vary independently, and between-plate variation was presumed to be normal on the log_e_-scale. The response variable was the difference in absorbance between 570 and the reference wavelength 630. Its residual error was defined as lognormal.

To provide an absolute scale for the effect, the E_MAX_ was defined on the logistic scale (constraining its value to (0,1), exclusive), and transformed to the observed scale as a multiple of the difference between the estimated response for DMSO controls, which were taken to represent the minimum possible response, and the E_MIN_. The E_MIN_ was constrained to be larger than the response for the DMSO controls such that the form of the dose-response relationship was forced to be negative (inhibitory). This specification follows (Long et al. 2025 – under review).

The prior information for all pharmacodynamic parameters and their variabilities was intended to be weak, simply constraining them to physically realistic values. The prior specification is available in the study code. Principles for specification of prior models for veterinary pharmacology applications were described by the author (Woodward, 2024). Goodness-of-fit of the completed models was assessed visually by comparing posterior predictions to the observations at the subject and plate level (**Figure 5**).

## Declarations

### Ethics approval and consent to participate

All sample collections derived from canine patients at Iowa State University (IACUC-21-250) and the University of Georgia (A2023 10-002-A1) were used under approved IACUC protocols. All biospecimen collections obtained from Purdue University were conducted under an approved IACUC protocol (#1111000124), and tissue or urine samples were shipped to ISU or UGA.

### Consent for publication

Not applicable.

### Availability of data and material

The stranded mRNA raw RNA-seq reads and snRNA-seq reads generated and analyzed in this study are available in the NCBI GEO (Accessions GSE306810 and GSE306811). Additionally, three lines had their RNA-seq files posted on the NIH ICDC portal (https://caninecommons.cancer.gov/#/study/ORGANOIDS01). The WGS raw files from this study are available in the Sequence Read Archive (NCBI-SRA BioProject PRJNA1312855).

RNA-seq and cytotoxicity scripts are available on GitHub (https://github.com/chris-zdyrski/TCC-Organoids). Scripts used for post-cram processing are also available on Github (https://github.com/Hexive-UN-03/cancer_code_release_2025) as well as to process to the cram stage WAGS was used up until the cram rule (https://github.com/jonahcullen/wags).

## Acknowledgments

This project was supported in part by Hannah Wickham, Dipak Kumar Sahoo, and Mohamed Elbadawy. We also thank Emily Rawlings and Maria E. Orbay-Cerrato for sample collection at the University of Georgia. We appreciate the collection of samples from Purdue University led by Deborah Knapp and Alexander Enstrom, as well as the Ethos Vet Hospital. We appreciate the timely processing of samples at the University of Georgia Histology Laboratory. Finally, we thank Fabrice Lucien-Matteoni and Numrah Fadra with their help for the bulk RNA-seq of three patients.

## Author contributions

C.Z., A.P. H.F.N., M.P.C., M.C., J.P., collected data and expanded organoid lines. E.F.D., A.P.W., H.H., and S.G.F. completed data analysis and software writing. M.P.C. and J.C. completed histological review. C.S. assisted in sample collection. C.Z., A.P., K.A., and J.P.M. wrote and edited the manuscript. K.A. and J.P.M. funded the project. All authors reviewed the manuscript.

## Declaration of generative AI in scientific writing

ChatGPT-4o (OpenAI, 2025) was used to improve the grammar and clarity of this manuscript. Its use was limited to enhancing readability. After using this tool, the authors reviewed and edited the content as needed and take full responsibility for the content of the publication.

## Supplemental Tables and Figures

### Patient metadata and detailed sample harvesting information

**Supplemental Table 1**: Additional patient metadata including diagnosis, patient location, and samples collected are listed.

### Microplate reader settings

**Supplemental Table 2**: Tecan Spark microplate reader settings used to assess cellular metabolic activity.

### Longitudinal sampling and organoid generation from one canine bladder cancer patient

**Supplemental Figure 1:**
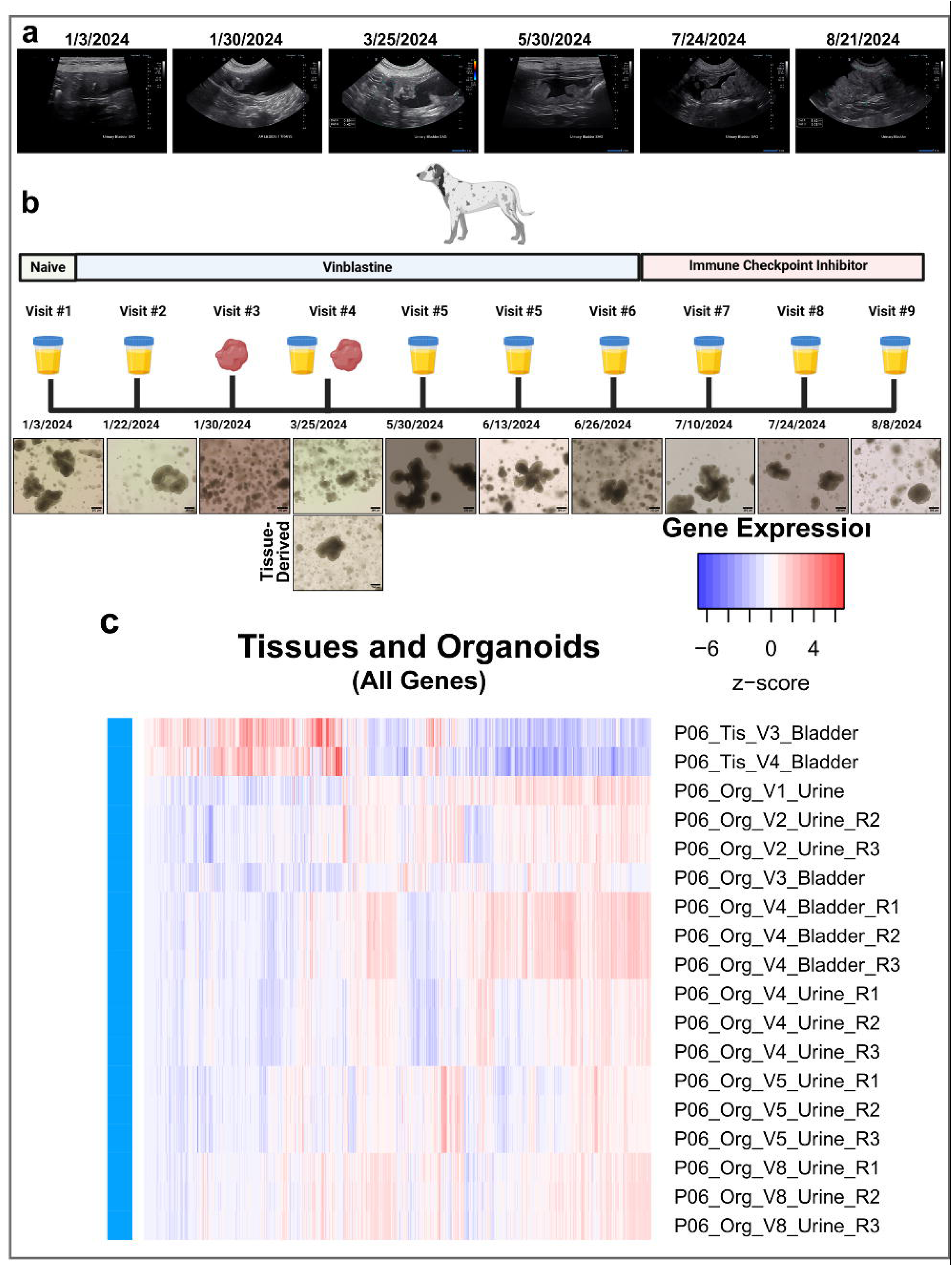
(**a**) Ultrasound imaging of the *in vivo* patient from visits throughout the growth of the tumor, from the same patient with which organoids were successfully derived, the tumor is visible within the bladder. (**b**) Types of samples collected over time and morphological photos for each one. Figure was made using Biorender.com. (**c**) RNA sequencing heatmap of all genes for longitudinal urine and tissue-derived organoid lines and tissue biopsies derived from P6.

**Supplemental Figure 2:**
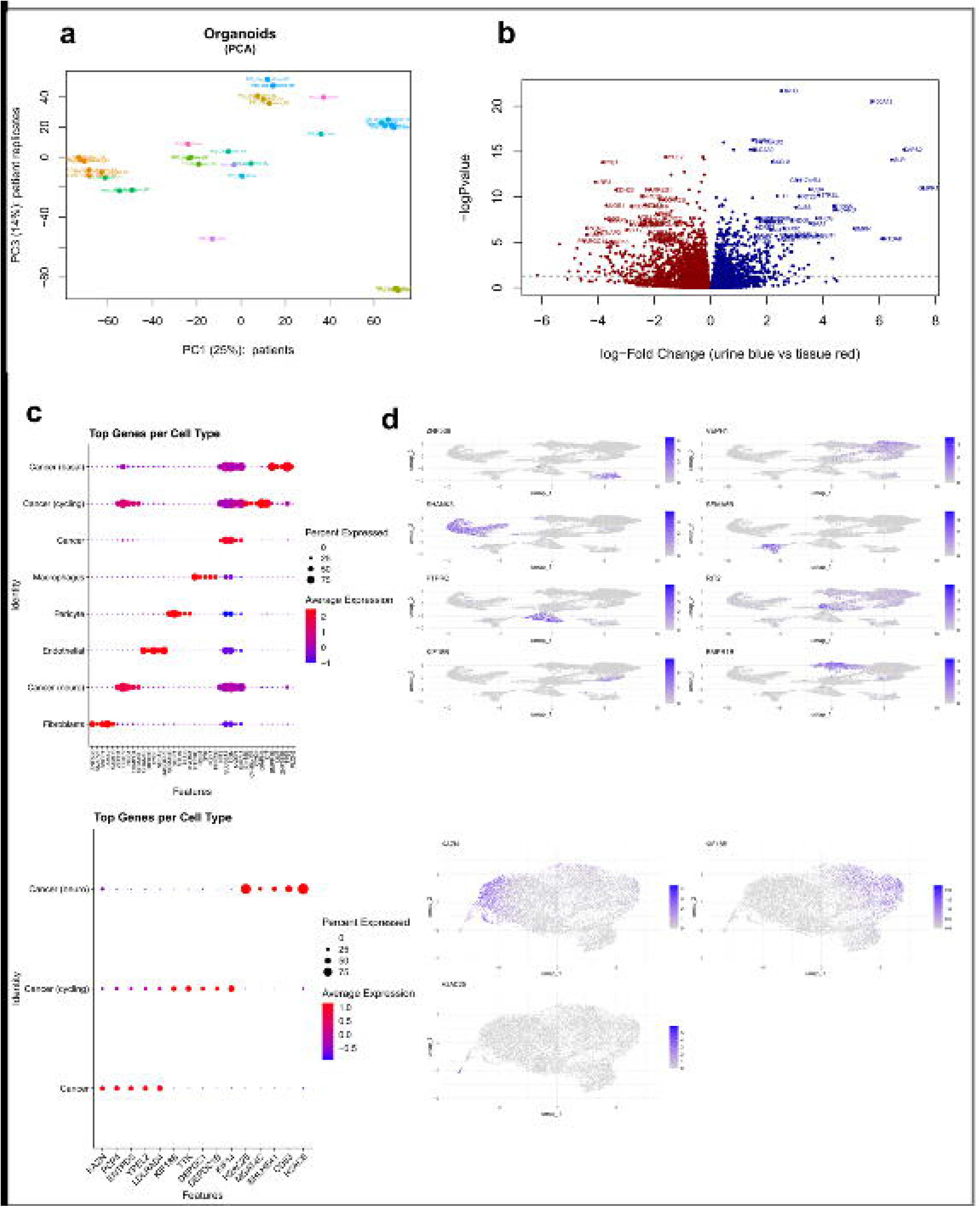
(**a**) PCA showing heterogeneity between transcriptomes of individual canine patient-derived UC organoid lines. Organoid lines derived from the same patients are depicted in the same color. (**b**) Volcano plot comparing the transcriptomes of all tissue-versus urine-derived organoids, where the dotted line represents significance. (**c**) Expression of the top 5 genes per cluster with the dot size depicting the percentage of cells in a class and dot color corresponding to the average expression level across all cells within a class (red = higher expression, blue = lower expression). (**d**) Selected genes from each cluster plotted against all other clusters.

